# Dietary restriction in the long-chain acyl-CoA dehydrogenase knockout mouse

**DOI:** 10.1101/2020.12.03.408591

**Authors:** Eugène F. Diekman, Michel van Weeghel, Mayte Suárez-Fariñas, Carmen Argmann, Pablo Ranea-Robles, Ronald J.A. Wanders, Gepke Visser, Ingeborg van der Made, Esther E. Creemers, Sander M. Houten

## Abstract

Patients with a disorder of mitochondrial long-chain fatty acid β-oxidation (FAO) have reduced fasting tolerance and may present with hypoketotic hypoglycemia, hepatomegaly, (cardio)myopathy and rhabdomyolysis. Patients should avoid a catabolic state because it increases reliance on FAO as energy source. It is currently unclear whether weight loss through a reduction of caloric intake is safe in patients with a FAO disorder. We used the long-chain acyl-CoA dehydrogenase knockout (LCAD KO) mouse model to study the impact of dietary restriction (DR) on the plasma metabolite profile and cardiac function. For this, LCAD KO and wild type (WT) mice were subjected to DR (70% of ad libitum chow intake) for 4 weeks and compared to ad libitum chow fed mice. We found that DR had a relatively small impact on the plasma metabolite profile of WT and LCAD KO mice. Echocardiography revealed a small decrease in left ventricular systolic function of LCAD KO mice, which was most noticeable after DR, but there was no evidence of DR-induced cardiac remodeling. Our results suggest that weight loss through DR does not have acute and detrimental consequences in a mouse model for FAO disorders.

## Introduction

Defects in mitochondrial long-chain fatty acid β-oxidation (FAO) compromise energy homeostasis in particular in catabolic states and lead to the accumulation of fatty acid derivatives such as acylcarnitines and triglycerides [1]. Patients with such a defect may develop hypoketotic hypoglycemia, hepatomegaly, cardiomyopathy and myopathy with rhabdomyolysis. These symptoms typically occur or may progress in a catabolic state induced by illness, fever, exercise or fasting. Under these conditions there is increased reliance on FAO for the production of ATP. Patients with a long-chain FAO disorder are treated with dietary intervention including restriction of long-chain fat intake and use of medium-chain triglycerides (MCT), triheptanoin and carbohydrates as alternative caloric sources [2]. In all FAO disorders (including medium-chain acyl-CoA dehydrogenase (MCAD) deficiency) patients are advised to prevent catabolic states. This can be achieved by emergency admissions in a hospital for glucose infusion during acute illness, but also by intake of MCT, carbohydrates or ketone bodies prior to exercise [3]. In some cases, late-evening meals and bedtime snacks are encouraged. Many countries have included FAO disorders in their newborn screening program in order to identify patients before they develop symptoms and improve disease outcome [4, 5].

In a natural history study of MCAD deficiency, it was noted that prepubertal children showed a tendency to be overweight. This was attributed to the dietary intervention or changes in feeding behavior [6]. Whether related to the deficiency and its treatment or not, overweight and obesity are common and may occur also in patients with a FAO defect. Given the health problems associated with being overweight or obese, patients with a FAO disorder may wish to lose weight through a diet and/or other lifestyle changes. Losing weight, however, is inevitably associated with a negative energy balance and catabolic state and therefore contraindicated in FAO disorders. Successful weight loss under supervision of a metabolic dietician was reported in two adult patients with a FAO defect [7].

The consequences of weight loss through dietary intervention in patients with a FAO disorder can be investigated in a model organism. Several mouse models with defect in long-chain FAO have been developed. We and others have shown that the long-chain acyl-CoA dehydrogenase (LCAD) KO mouse presents with phenotypes that resemble some aspects FAO disorders such as very long-chain acyl-CoA dehydrogenase (VLCAD) deficiency. These phenotypes include increased C12- and C14-acylcarnitines, fatty liver, fasting-induced hypoketotic hypoglycemia and cardiac hypertrophy [8-10]. It is, however, unknown what the impact of dietary restriction (DR) is on LCAD KO mice. We hypothesized that DR increases FAO and therefore can aggravate the phenotype of LCAD KO mice.

## Materials and methods

### Animals

Acadl^-/-^ mice (B6.129S6-Acadl^tm1Uab^/Mmmh [8]) on a pure C57BL/6N background were obtained from Mutant Mouse Regional Resource Centers (http://www.mmrrc.org/). Colonies were maintained in by regular crossing with C57BL/6N mice in order prevent genetic drift. Experimental cohorts (all male animals) were generated by homozygote breeding pairs of mice after 16 generations of backcrossing and 4 or 5 generations of line intercrosses. The LCAD allele was genotyped using PCR in combination with acylcarnitine analysis on bloodspots with C14:1/C2 ratio being predictive for the LCAD KO genotype [9]. Mice were housed at 21 ± 1°C, 40-50% humidity, on a 12-h light-dark cycle, with ad libitum access to water and a standard rodent diet (chow; CRM (E), Special Diets Services) until the mice were ∼8 weeks of age. At that moment, mice were housed individually, body weights were measured and 50µl blood was drawn via the vena saphena for measurement of glucose (with glucometer), ketones, acylcarnitines, lactate and pyruvate. After this, one group (6 WT and 5 KO mice) continued on the standard ad libitum feeding of chow diet. A second group (6 WT and 6 KO mice) was subjected to dietary restriction (DR) receiving 70% of the chow intake in the parallel fed control group (2.8 to 3.0 g of chow per day). During the 4 week study period, body weight was measured biweekly. After two weeks, a 50µl blood was drawn via the vena saphena for measurement of glucose (with glucometer), ketones, acylcarnitines, creatine kinase (CK), lactate and pyruvate. Echocardiography was performed at baseline, and after 4 weeks of dietary intervention (before necropsy). For the necropsy, mice were first anesthetized with pentobarbital (100mg/kg via intraperitoneal injection) and then exsanguinated via the vena cava inferior. Blood was collected for perchloric acid extraction or the preparation of EDTA plasma and organs were weighed and snap frozen in liquid nitrogen and stored at -80°C for future analyses. The blood and plasma was used for measurement of glucose, β-hydroxybutyrate, lactate, pyruvate, amino acids, glycerol, free fatty acids, triglycerides, acylcarnitines and CK. Data and samples for this experiment were collected in 2 animal cohorts; cohort 1 in October 2011 (4 WT chow, 5 WT DR, 3 KO chow and 3 KO DR) and cohort 2 in February 2012 (2 WT chow, 1 WT DR, 2 KO chow and 3 KO DR). All experiments were approved by the institutional review board for animal experiments at the Academic Medical Center (Amsterdam, The Netherlands) and comply with the National Institutes of Health guide for the care and use of Laboratory animals (NIH Publications No. 8023, revised 1978).

### Clinical chemistry measurements

Blood for measurement of lactate, pyruvate and ketones was treated immediately after withdrawal with an equal volume of 1M perchloric acid and subsequently stored at -20°C. The remainder of the blood was used for plasma preparation. Plasma was stored at -20°C. Blood glucose was determined using a Contour glucometer (Bayer, Mijdrecht, The Netherlands). Blood lactate, pyruvate, β-hydroxybutyrate and plasma amino acids and acylcarnitines were measured using tandem mass spectrometry. Glycerol (Instruchemie, 2913), free fatty acids (Wako, 434-91795), triglycerides (Human Gesellschaft, 10724) and creatine kinase were measured in plasma using standard enzymatic assays as described [9, 11].

### Echocardiography

Left ventricular dimensions and systolic function were determined by transthoracic 2D echocardiography using a Vevo 770 Ultrasound (Visual Sonics) equipped with a 30-MHz linear array transducer [12]. Mice were sedated on 4% isoflurane and anesthesia was maintained by a mixture of O_2_ and 2.5% isoflurane. M-mode tracings in parasternal short axis view at the height of the papillary muscle were used to measure left ventricular (LV) anterior wall (AW) thickness, LV posterior wall (PW) thickness, LV internal diameter (LVID) at end-systole (s) and end-diastole (d). Ejection fraction (EF) and fractional shortening (FS) were calculated from the LVID as described [13].

### Immunoblot analysis

Immunoblotting of mouse heart homogenates was performed as described in an earlier study using the same primary antibodies [14].

### Statistics

Statistical analysis was performed using Graphpad Prism 6 as indicated in the figure and table legends. Data are displayed as the mean ± confidence interval as indicated in the figure legend or table. A linear mixed effect model was used to model the echocardiography data using R. Time, diet and genotype were fixed factors and the individual animals were a random factor. Statistical significance is indicated as follows: + P<0.1; * P < 0.05, ** P <0.01, and *** P<0.001.

## Results

### Dietary restriction in the LCAD KO mouse

At the start of the study (∼8 weeks of age) mice were single housed. Half of the study cohort continued ad libitum feeding (chow), whereas the other half was subjected to dietary restriction (DR) (Figure 1A). Basal body weights of LCAD KO mice were slightly higher when compared to the WT mice (27.9 ± 0.5 versus 26.1 ± 0.4, respectively, P=0.0126). In the first days of the study all mice lost body weight likely due to the stress associated with single housing. Body weight loss was more pronounced in the DR group. After 1 week, body weights had recovered in the chow groups. Body weights in the DR groups slowly decreased during the study period (Figure 1B). There was no difference in body weight change between WT and LCAD KO mice in chow and DR groups at the end of the study period (Table S1, S2). DR decreased heart, liver and adipose weight in WT and LCAD KO mice. Heart weight was increased in the LCAD KO, which was most pronounced in the DR group upon normalization for body weight (Figure 1C).

**Figure 1.**
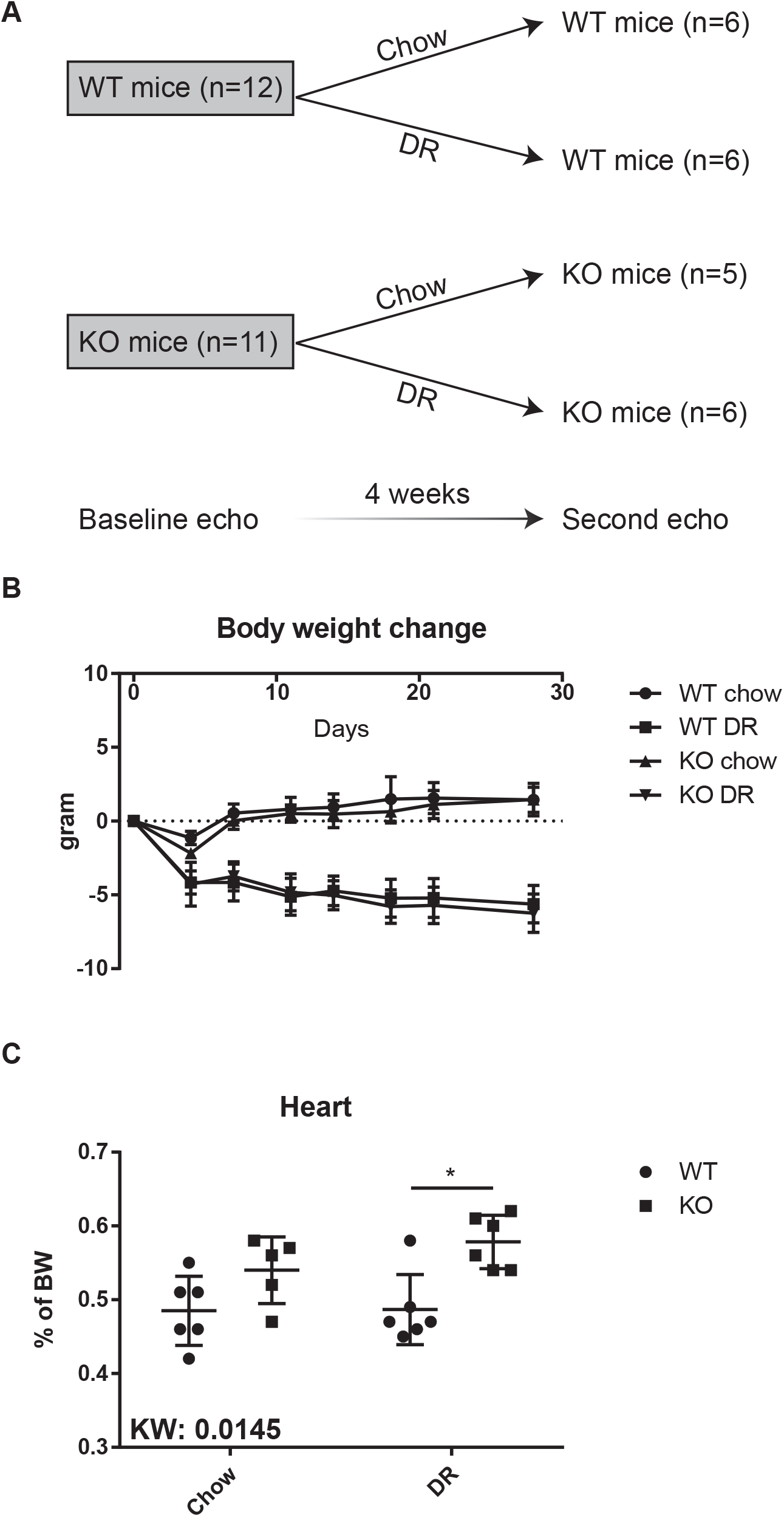
Dietary restriction in the LCAD KO mouse. (A) Schematic representation of the study design. (B) Body weight change was calculated as the difference of the body weight at the indicated time point and the body weight at the baseline measurement (day 0). Mean ± SD. (C) Heart weight normalized as percentage of body weight. The result of a Kruskal-Wallis and Dunn’s multiple comparisons test are displayed. Mean ± SD.

### Biochemical characterization of LCAD KO mice during dietary restriction

LCAD KO mice develop hypoketotic hypoglycemia upon fasting [9]. We measured blood glucose, lactate, pyruvate and ketone bodies after 2 and 4 weeks of dietary intervention. Although quite variable, blood glucose was lower in mice of the DR groups with the lowest values observed in LCAD KO mice (Figure 2A). Levels of the ketone body β-hydroxybutyrate were also quite variable, but generally higher in the DR groups (Figure 2A). Unexpectedly, the mice with the highest plasma β-hydroxybutyrate concentration were observed in the LCAD KO DR group. Blood lactate and pyruvate levels were not different between the various groups (Table S1, S2 and not shown).

**Figure 2.**
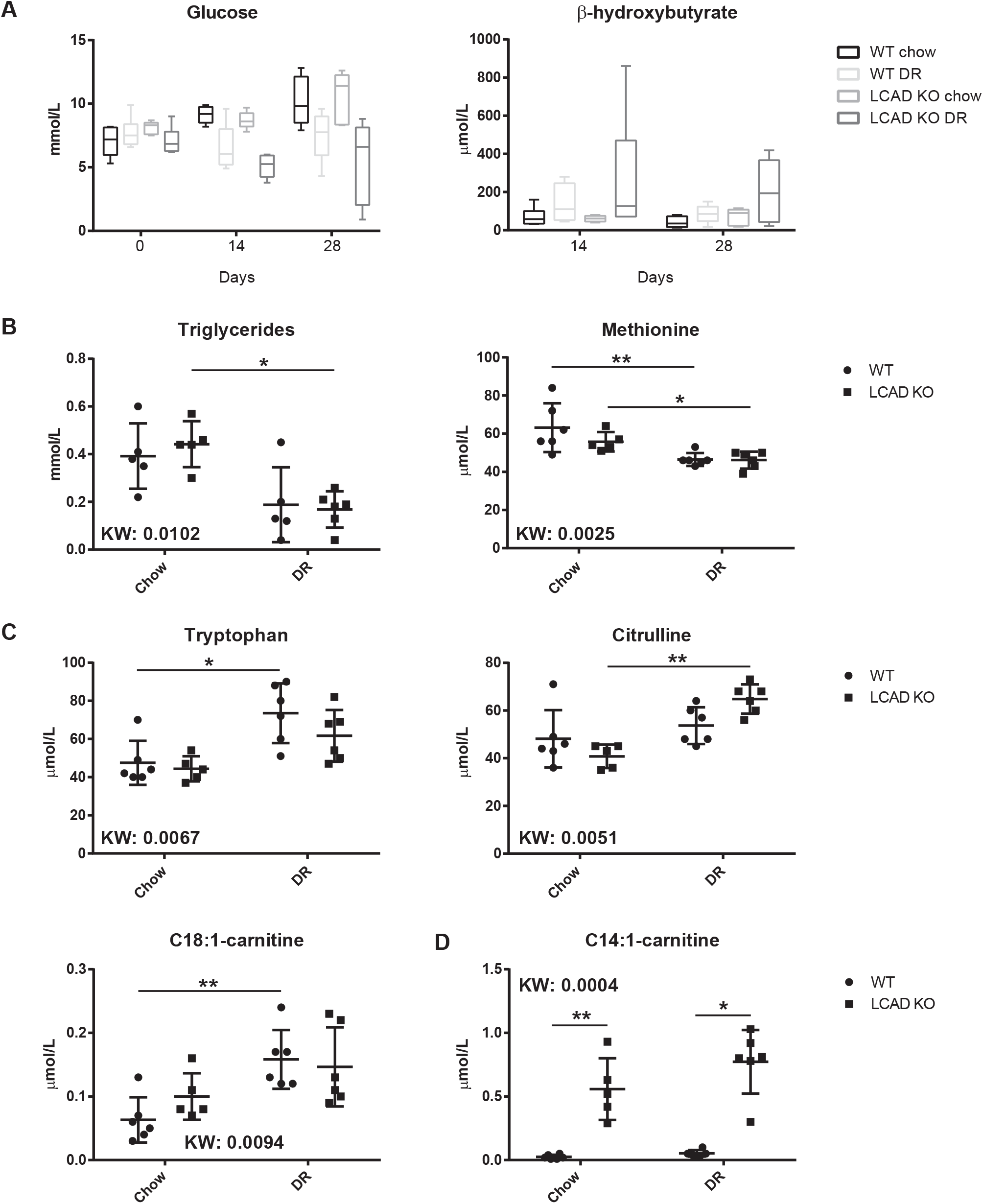
Biochemical characterization of LCAD KO mice during dietary restriction. (A) Blood glucose and β-hydroxybutyrate concentration at different time points during the experiment. The results of the repeated measures two-way ANOVA (as % of total variation explained) for glucose are: interaction 15.54% (P=0.0142), time point 4.59% (not significant), group 34.54% (P<0.0001) and subjects (matching) 14.04% (not significant). A repeated measures two-way ANOVA could not be performed for β-hydroxybutyrate because of missing values for some subjects. A two-way ANOVA showed a significant group effect 24.62% (P=0.0162). Box and whiskers indicating 5-95 percentile. (B) Plasma concentration of triglycerides and methionine. (C) Plasma concentration of tryptophan, citrulline and C18:1-carnitine. (D) Plasma concentration of C14:1-carnitine. For B, C and D panels, the result of a Kruskal-Wallis and Dunn’s multiple comparisons test are displayed. Mean ± SD.

A more detailed characterization of plasma metabolites including amino acids and acylcarnitines was performed at the end of the study (Table S1, S2). Triglycerides and methionine were decreased by DR (Figure 2B). Tryptophan, citrulline and several C16- and C18-acylcarnitines were increased by DR (Figure 2C). The DR-dependent increase in citrulline was most pronounced in LCAD KO mice, whereas the increase in C16- and C18-acylcarnitines was most marked in WT animals. As expected, the C12- and C14-acylcarnitines were mostly affected by genotype (Figure 2D, S1). Free carnitine, acetylcarnitine and C4OH-carnitine were only affected by genotype in DR mice (Figure S1). Plasma CK levels were within the normal range indicating that LCAD KO mice do not develop muscle damage during DR (not shown). Taken together, DR had a relatively small impact on the plasma metabolite profile of LCAD KO mice.

### Cardiac function in LCAD KO mice before and after dietary restriction

We assessed the effects of DR on left ventricular (LV) systolic function of WT and LCAD KO mice using echocardiography. For this, we measured anterior wall thicknesses (AW), left ventricular internal dimension (LVID) and posterior wall (PW) thicknesses at diastole (d) and systole (s) at baseline (0 weeks) and after 4 weeks of ad libitum feeding or DR (Table S3, S4, and S5). At baseline, LCAD KO mice had an increased AWd and AWs, but no other significant differences were noted. Next, we used a linear mixed effect model to investigate the effects of time (week 0 versus week 4), diet (chow versus DR), genotype (WT vs KO) and their interactions (Figure 3A). Time affected heart rate (HR), body weight (BW), PWd, PWs and AWs, diet affected HR, BW and AWd, and genotype affected BW, PWd, PWs, AWd, AWs, EF and FS.

**Figure 3.**
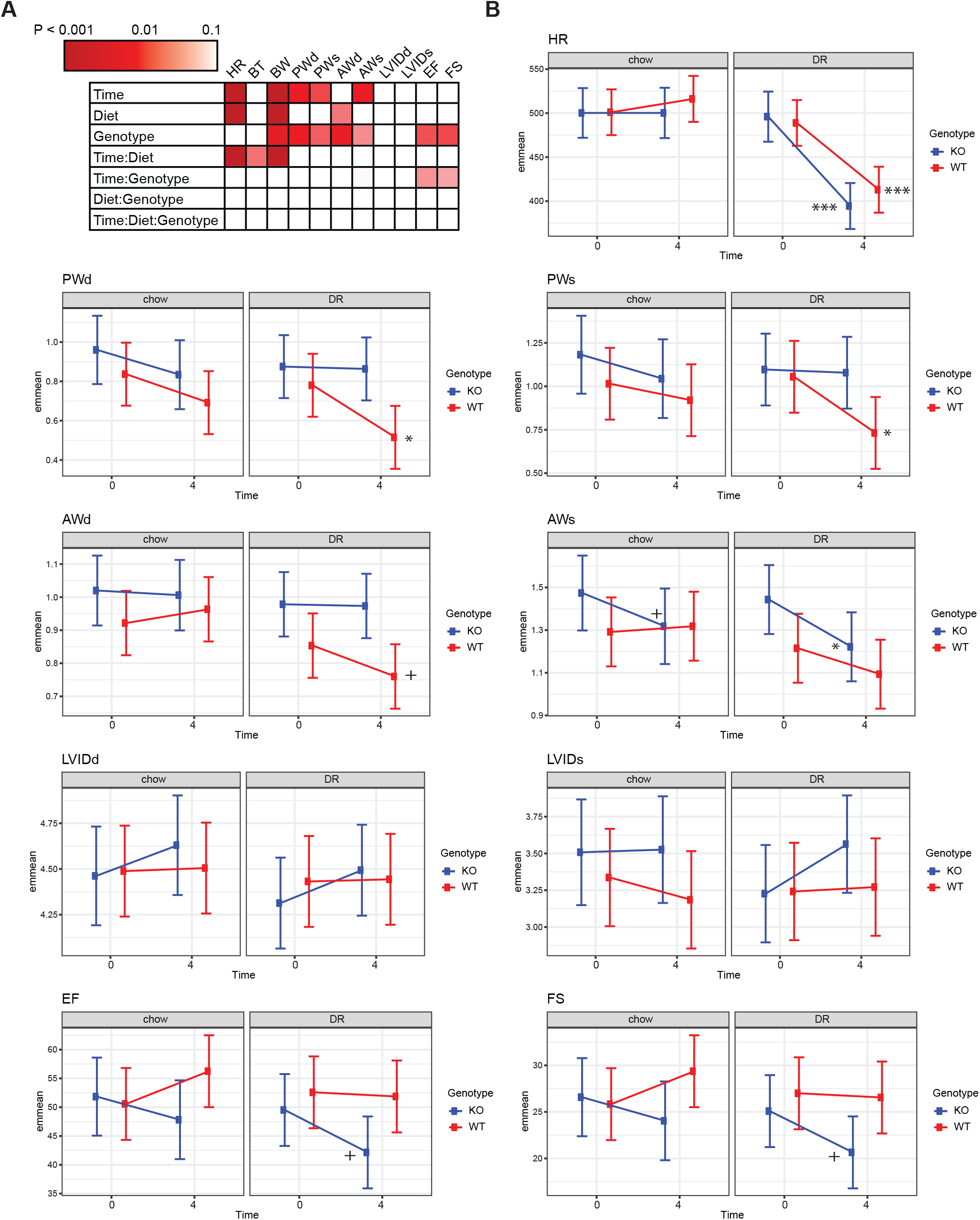
Cardiac function in LCAD KO mice before and after dietary restriction. (A) Heat map displaying P values for the fixed variables time, diet and genotype and their interaction on the measured echocardiography variables in the linear mixed effect model. (B) Individual graphs of the measured echocardiography variables. The level of significance for the diet-induced changes is indicated in the graphs, for WT to the right of time point 4, for LCAD KO to the left of time point 4. Error bars indicate the upper and lower limit of the 95% confidence interval.

Time and diet significantly interacted for HR and BW (Figure 3A), which is explained by the pronounced decrease in HR and BW upon DR (Figure 3B and S2). The genotype effect on BW is explained by the increased BW in the LCAD KO group at baseline. PWd, PWs, AWd and AWs were overall higher in LCAD KO animals (Figure 3B), which is consistent with cardiac hypertrophy. PWd, PWs and AWd all decreased in WT animals upon DR (Figure 3B), which may reflect loss of cardiac mass due to DR. This decrease was not observed in LCAD KO mice and therefore differences in PWd, PWs and AWd between with the WT and LCAD KO mice were most pronounced after DR (Figure 3B). AWs decreased in LCAD KO animals in chow and DR condition (Figure 3B). Time and genotype interacted for EF and FS (P<0.1, Figure 3A). EF and FS decreased with time in LCAD KO under chow and DR conditions, but this decrease was most pronounced in the DR condition (Figure 3B). In WT mice, EF and FS did not change much explaining the time;genotype interaction.

In order to assess if DR caused remodeling of the hypertrophied LCAD KO heart, we studied the protein expression of brain natriuretic peptide (BNP; NPPB), myosin heavy chain 7 (MYH7, β-myosin heavy chain), myosin light chain 1 (MYL1) and α-SMA (ACTA2, α-smooth muscle actin) in the mice from the first cohort (Figure 4). BNP and MYH7 are well-established markers of cardiac remodeling during pathological hypertrophy. The role of MYL1 in cardiac remodeling is unknown, but we have noted that its expression is decreased in LCAD KO hearts [14]. α-SMA is a marker to detect the presence of myofibroblasts in cardiac fibrosis. Expression of BNP was overall comparable between samples, but one LCAD KO on DR had notable higher expression levels. This mouse had the lowest EF and FS at baseline and after 4 weeks of DR arguing against DR as a cause of the increased BNP. Expression of MYH7 and MYL1 was induced by DR, but the increase was less pronounced in LCAD KO samples. Expression of α- SMA was higher in the DR samples, but there was no effect of genotype. These data suggest that cardiac hypertrophy in the LCAD KO mouse is non-pathological under both the ad libitum feeding and DR conditions.

**Figure 4.**
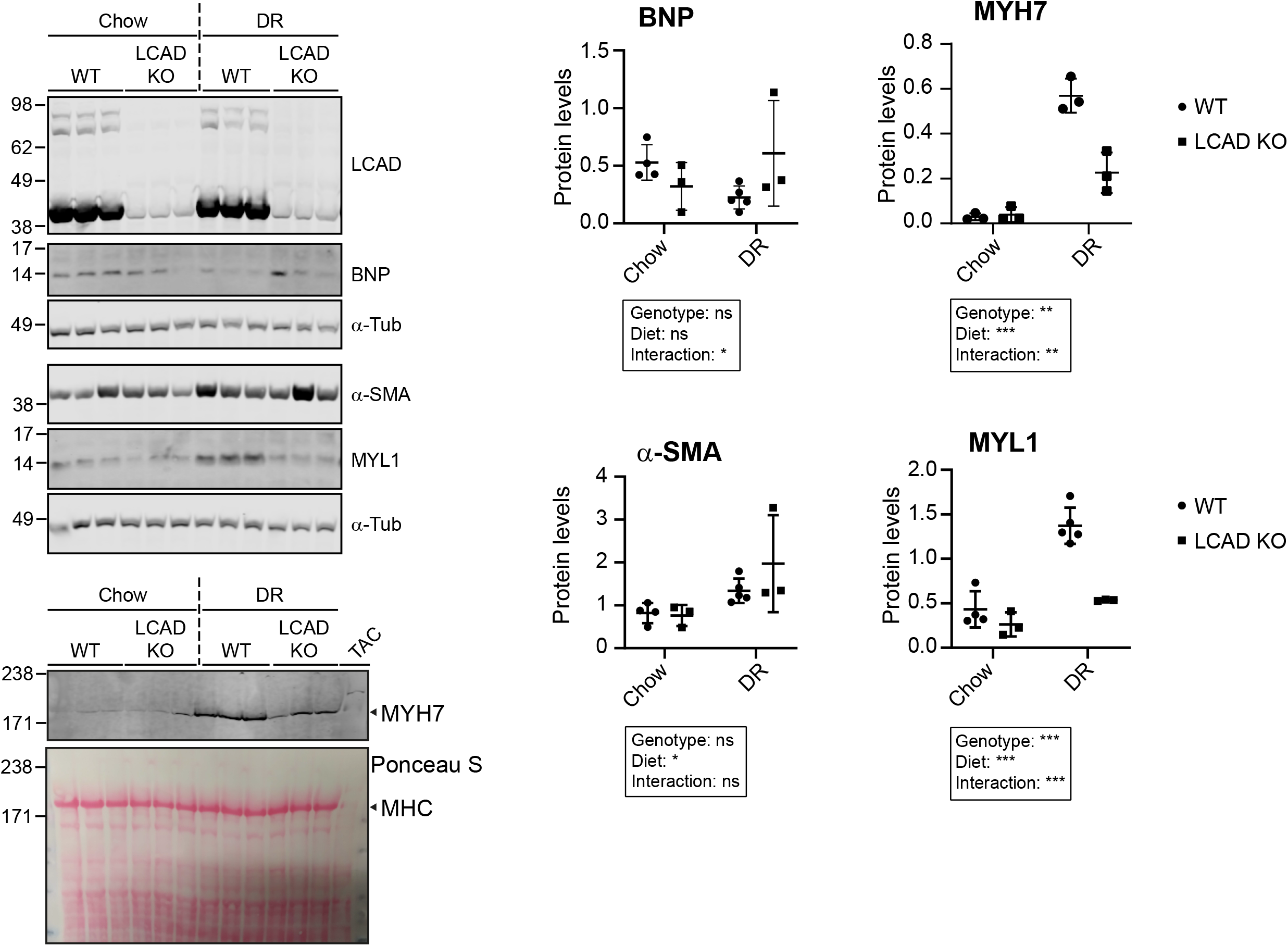
Immunoblots of markers for cardiac remodeling in WT and LCAD KO mice after 4 weeks of chow ad libitum feeding or DR. Only mice from cohort 1 were used for this experiment (4 WT chow, 5 WT DR, 3 KO chow and 3 KO DR). The last lane of the MYH7 immunoblot includes a sample from a mouse heart after transverse aortic constriction (TAC) as positive control. For each separate blot, the α-tubulin loading control is displayed. For the MYH7 blot, the Ponceau S stain is shown and the abundant myosin heavy chain (MHC) is indicated. The positions of the molecular weight markers are indicated in kDa. Quantification of relative protein levels is provided, and the results of the two-way ANOVA are displayed.

## Discussion

Dietary interventions in patients with FAO disorders aim to prevent metabolic decompensation as a consequence of a catabolic state. This may pose a problem for those patients who wish to lose weight because they are overweight or obese, as this is inevitably associated with a catabolic state and therefore risky. Here we investigated the consequences of long term DR in the LCAD KO, a model for long-chain FAO disorders. We focused on the heart as patients with long-chain FAO disorders are at risk of developing cardiomyopathy and/or arrhythmias, and LCAD KO mice are known to have cardiac hypertrophy.

We also provided a detailed characterization of plasma metabolites. LCAD KO mice on DR were expected to display hypoketotic hypoglycemia similar to overnight fasted LCAD KO mice. Although highly variable, the lowest blood glucose values were indeed observed in LCAD KO mice on DR. Unexpectedly, LCAD KO on DR developed moderate ketosis and displayed the highest β-hydroxybutyrate concentrations observed in this study. We speculate that this reflects their lower fasting tolerance, which is exemplified by a rapid onset (within 4 hours) of hypoglycemia during fasting due to increased glucose requirement [9]. Although circulating ketone bodies are decreased in overnight fasted LCAD KO mice, they are not absent and clearly higher than the levels in the fed and postabsorptive state [9]. This suggests that residual hepatic FAO (via VLCAD) as well as oxidation of ketogenic amino acids such lysine and leucine allows for some ketogenesis in the LCAD KO mouse.

Cardiac function of LCAD KO mice has been compared to WT controls using echocardiography and magnetic resonance imaging (MRI) [10, 15, 16]. Echocardiography has revealed that interventricular septal and LV PW thicknesses were increased in LCAD KO mice, without any apparent changes in LV systolic function as assessed by EF and FS [16]. Female sex and phytoestrogens from diet were found to be protective against hypertrophy [16]. Similarly, in the MRI studies, cardiac function was largely maintained in LCAD KO mice in the fed state [10, 15, 17]. Fasting, however, caused an impaired LV performance in LCAD KO mice including an increase in end systolic volume, and decreases in EF, stroke volume, cardiac output, peak ejection rate and peak late filling rate [10, 15]. The cardiac phenotype of LCAD KO mice is reminiscent of the subclinical effects observed in adults with a FAO disorder, which included a moderately increased LV mass, increased LV torsion during ejection and largely preserved EF [18]. Here, we used echocardiography to monitor the effects of DR on LV systolic function in WT and LCAD KO mice. AW thicknesses were increased in LCAD KO mice at baseline. After DR, PW thicknesses had decreased in WT, but not LCAD KO mice. This result is reminiscent of our previous study in which LV mass decreased in WT mice upon overnight fasting, an effect that was not observed in LCAD KO mice [10]. No differences between cardiac function of WT and LCAD KO mice were noted at baseline analysis, but EF and FS were affected by genotype (P<0.05) with near significant time:genotype interactions (P<0.1). Although the lowest values were found in the DR group, cardiac function also decreased in the chow group. The DR-induced changes were near significant in the LCAD KO group (P<0.1), but immunoblotting for BNP and MYH7 revealed no evidence of pathological cardiac remodeling. It is unclear why there was a general decline in cardiac function in the LCAD KO group. One possible explanation may be the single housing, which can be stressful to social animals such as mice.

In our experiment, DR led to a negative energy balance characterized by an initial rapid decline in body weight followed by a slower body weight decrease over time. Surprisingly, DR did not appear to lead to a major impairment in cardiac function or elevated CK. We speculate that mice minimize the consequences of decreased caloric availability by reducing energy expenditure. Indeed, mice are known to exhibit a robust cardiovascular response to both acute and chronic negative energy balance [19, 20]. This is consistent with the pronounced decrease in heart rate in WT and LCAD KO mice upon DR (Figure 3B). Previously, we have demonstrated that food withdrawal can lower energy expenditure and induce inactivity in the LCAD KO mouse model [11]. This ability of mice to adapt to a negative energy balance differs from humans and may make translation of some findings challenging.

There are additional limitations in our study that may have limited to impact of DR in the LCAD KO mouse model. Firstly, due to overlap of LCAD and VLCAD function [21], LCAD KO mice have a relatively mild phenotype, which includes fasting-induced hypoketotic hypoglycemia, cardiac hypertrophy and increased liver weight [8-10]. In contrast to the human long-chain FAO disorders, these phenotypes generally do not progress to a life-threatening condition. In addition, skeletal muscle myopathy has not been observed in LCAD KO mice upon food withdrawal [11]. Secondly, our study may have been underpowered to detect relevant differences. Indeed, some of the highlighted effects missed the traditional P value cutoff for statistical significance (0.05). Regardless, the detected effects in EF and FS are rather small and may have limited biological relevance. Lastly, the mice on DR received the same chow as the ad libitum fed animals. Chow is formulated based on ad libitum intake, and restriction of this diet may cause nutritional deficiencies of vitamins, minerals or other essential nutrients [22]. We measured plasma amino acids and found that of the essential amino acids only the concentrations of methionine and tryptophan were changed by DR, but only methionine was decreased. This may be consistent with methionine and cysteine being slightly below recommended intake due to the DR [23].

In summary, 4 weeks of DR in LCAD KO mice does not lead to pronounced worsening of existing hypertrophic cardiomyopathy. Cardiac hypertrophy overall remained mild with no signs of pathological remodeling. Although cardiac function was slightly decreased in LCAD KO mice, DR contributed little to this decrease. Our findings suggest that mice with a FAO defect can tolerate longer periods with a negative energy balance. Given the subclinical effects of a FAO defect on the adult heart [18], weight loss in patients should be done gradual and under supervision of medical specialists with appropriate monitoring of cardiac function and plasma CK.

## Supporting information

All supplemental materials

## Acknowledgements

Research reported in this publication was supported by the National Institute of Diabetes and Digestive and Kidney Diseases of the National Institutes of Health under Award Number R01DK113172. The content is solely the responsibility of the authors and does not necessarily represent the official views of the National Institutes of Health. This work was also supported by the Netherlands Organization for Scientific Research (VIDI-grant No. 016.086.336 to SMH), ZonMW (dossier 200320006), Metakids (www.metakids.nl).

## References

[1] S.M. Houten, S. Violante, F.V. Ventura, R.J. Wanders, The biochemistry and physiology of mitochondrial fatty acid beta-oxidation and its genetic disorders, Annu Rev Physiol, 78 (2016) 23–44.

[2] M.B. Gillingham, S.B. Heitner, J. Martin, S. Rose, A. Goldstein, A.H. El-Gharbawy, S. Deward, M.R. Lasarev, J. Pollaro, J.P. DeLany, L.J. Burchill, B. Goodpaster, J. Shoemaker, D. Matern, C.O. Harding, J. Vockley, Triheptanoin versus trioctanoin for long-chain fatty acid oxidation disorders: a double blinded, randomized controlled trial, J Inherit Metab Dis, 40 (2017) 831–843.

[3] J.C. Bleeker, G. Visser, K. Clarke, S. Ferdinandusse, F.H. de Haan, R.H. Houtkooper, I.J. L. I.L. Kok, M. Langeveld, W.L. van der Pol, M.G.M. de Sain-van der Velden, A. Sibeijn-Kuiper, T. Takken, R.J.A. Wanders, M. van Weeghel, F.A. Wijburg, L.H. van der Woude, R.C.I. Wust, P.J. Cox, J.A.L. Jeneson, Nutritional ketosis improves exercise metabolism in patients with very long-chain acyl-CoA dehydrogenase deficiency, J Inherit Metab Dis, 43 (2020) 787–799.

[4] V. Rovelli, F. Manzoni, K. Viau, M. Pasquali, N. Longo, Clinical and biochemical outcome of patients with very long-chain acyl-CoA dehydrogenase deficiency, Mol Genet Metab, 127 (2019) 64–73.

[5] J.C. Bleeker, I.L. Kok, S. Ferdinandusse, W.L. van der Pol, I. Cuppen, A.M. Bosch, M. Langeveld, T.G.J. Derks, M. Williams, M. de Vries, M.F. Mulder, E.R. Gozalbo, M.G.M. de Sain-van der Velden, A.J. Rennings, P. Schielen, E. Dekkers, R.H. Houtkooper, H.R. Waterham, M.L. Pras-Raves, R.J.A. Wanders, P.M. van Hasselt, M. Schoenmakers, F.A. Wijburg, G. Visser, Impact of newborn screening for very-long-chain acyl-CoA dehydrogenase deficiency on genetic, enzymatic, and clinical outcomes, J Inherit Metab Dis, 42 (2019) 414–423.

[6] T.G. Derks, D.J. Reijngoud, H.R. Waterham, W.J. Gerver, M.P. van den Berg, P.J. Sauer, G.P. Smit, The natural history of medium-chain acyl CoA dehydrogenase deficiency in the Netherlands: clinical presentation and outcome, J. Pediatr., 148 (2006) 665–670.

[7] H. Zweers, C. Timmer, E. Rasmussen, M. den Heijer, H. de Valk, Successful weight loss in two adult patients diagnosed with late-onset long-chain Fatty Acid oxidation defect, JIMD Rep, 6 (2012) 127–129.

[8] D.M. Kurtz, P. Rinaldo, W.J. Rhead, L. Tian, D.S. Millington, J. Vockley, D.A. Hamm, A.E. Brix, J.R. Lindsey, C.A. Pinkert, W.E. O’Brien, P.A. Wood, Targeted disruption of mouse longchain acyl-CoA dehydrogenase gene reveals crucial roles for fatty acid oxidation, Proc. Natl. Acad. Sci. USA, 95 (1998) 15592–15597.

[9] S.M. Houten, H. Herrema, H. te Brinke, S. Denis, J.P. Ruiter, T.H. van Dijk, C.A. Argmann, R. Ottenhoff, M. Muller, A.K. Groen, F. Kuipers, D.J. Reijngoud, R.J. Wanders, Impaired amino acid metabolism contributes to fasting-induced hypoglycemia in fatty acid oxidation defects, Hum. Mol. Genet., 22 (2013) 5249–5261.

[10] A.J. Bakermans, T.R. Geraedts, M. van Weeghel, S. Denis, M.J. Ferraz, J.M. Aerts, J. Aten, K. Nicolay, S.M. Houten, J.J. Prompers, Fasting-induced myocardial lipid accumulation in long-chain acyl-CoA dehydrogenase knock-out mice is accompanied by impaired left ventricular function, Circ. Cardiovasc. Imaging, 4 (2011) 558–565.

[11] E.F. Diekman, M. van Weeghel, R.J. Wanders, G. Visser, S.M. Houten, Food withdrawal lowers energy expenditure and induces inactivity in long-chain fatty acid oxidation-deficient mouse models, FASEB J., 28 (2014) 2891–2900.

[12] W.J. Wijnen, I. van der Made, S. van den Oever, M. Hiller, B.A. de Boer, D.I. Picavet, I.A. Chatzispyrou, R.H. Houtkooper, A.J. Tijsen, J. Hagoort, H. van Veen, V. Everts, J.M. Ruijter, Y.M. Pinto, E.E. Creemers, Cardiomyocyte-specific miRNA-30c over-expression causes dilated cardiomyopathy, PloS one, 9 (2014) e96290.

[13] S. Gao, D. Ho, D.E. Vatner, S.F. Vatner, Echocardiography in Mice, Current protocols in mouse biology, 1 (2011) 71–83.

[14] P. Ranea-Robles, C. Yu, N. van Vlies, F.M. Vaz, S.M. Houten, Slc22a5 haploinsufficiency does not aggravate the phenotype of the long-chain acyl-CoA dehydrogenase KO mouse, J Inherit Metab Dis, 43 (2020) 486–495.

[15] A.J. Bakermans, M.S. Dodd, K. Nicolay, J.J. Prompers, D.J. Tyler, S.M. Houten, Myocardial energy shortage and unmet anaplerotic needs in the fasted long-chain acyl-CoA dehydrogenase knockout mouse, Cardiovasc. Res., 100 (2013) 441–449.

[16] K.B. Cox, J. Liu, L. Tian, S. Barnes, Q. Yang, P.A. Wood, Cardiac hypertrophy in mice with long-chain acyl-CoA dehydrogenase or very long-chain acyl-CoA dehydrogenase deficiency, Lab. Invest., 89 (2009) 1348–1354.

[17] A.J. Bakermans, M. van Weeghel, S. Denis, K. Nicolay, J.J. Prompers, S.M. Houten, Carnitine supplementation attenuates myocardial lipid accumulation in long-chain acyl-CoA dehydrogenase knockout mice, J. Inherit. Metab. Dis., 36 (2013) 973–981.

[18] S.J.G. Knottnerus, J.C. Bleeker, S. Ferdinandusse, R.H. Houtkooper, M. Langeveld, A.J. Nederveen, G.J. Strijkers, G. Visser, R.J.A. Wanders, F.A. Wijburg, S.M. Boekholdt, A.J. Bakermans, Subclinical effects of long-chain fatty acid beta-oxidation deficiency on the adult heart: A case-control magnetic resonance study, J Inherit Metab Dis, 43 (2020) 969–980.

[19] T.L. Jensen, M.K. Kiersgaard, D.B. Sorensen, L.F. Mikkelsen, Fasting of mice: a review, Lab Anim, 47 (2013) 225–240.

[20] T.D. Williams, J.B. Chambers, R.P. Henderson, M.E. Rashotte, J.M. Overton, Cardiovascular responses to caloric restriction and thermoneutrality in C57BL/6J mice, Am J Physiol Regul Integr Comp Physiol, 282 (2002) R1459–1467.

[21] M. Chegary, H. te Brinke, J.P. Ruiter, F.A. Wijburg, M.S. Stoll, P.E. Minkler, M. van Weeghel, H. Schulz, C.L. Hoppel, R.J. Wanders, S.M. Houten, Mitochondrial long chain fatty acid beta-oxidation in man and mouse, Biochim. Biophys. Acta, 1791 (2009) 806–815.

[22] T.D. Pugh, R.G. Klopp, R. Weindruch, Controlling caloric consumption: protocols for rodents and rhesus monkeys, Neurobiol Aging, 20 (1999) 157–165.

[23] National Research Council, Nutrient Requirements of Laboratory Animals, Fourth Revised Edition ed., The National Academies Press, Place Published, 1995.

