## Supplementary material for "Dietary restriction in the long-chain acyl-CoA dehydrogenase knockout mouse": All supplemental materials

**Supplemental figure legends**

Figure S1. Biochemical characterization of LCAD KO mice during dietary restriction. Plasma concentration of C12-carnitine, free carnitine, C2-carnitine and C4OH-carnitine. The result of a Kruskal-Wallis and Dunn's multiple comparisons test are displayed. Mean ± SD.

Figure S2. Cardiac function in LCAD KO mice before and after dietary restriction. Individual graphs of body temperature (bodytemp) and body weight (BW). The level of significance for the diet-induced changes is indicated in the graphs, for WT to the right of time point 4, for LCAD KO to the left of time point 4. Error bars indicate the upper and lower limit of the 95% confidence interval.

### Supplementary tables

**Table S1.** Analyzed variables in WT and LCAD KO mice after 4 weeks of ad libitum chow or dietary restriction. The effect of the dietary intervention is calculated in a post hoc test.

| | KW | Chow | | post | Dietary restriction | | post |
|---|---|---|---|---|---|---|---|
| Parameters | Sig. | WT | LCAD KO | Sig. | WT | LCAD KO | Sig. |
| Body weight (g) | *** | 27.7 ± 1.3 | 29.5 ± 2.6 | ns | 20.2 ± 1.6 | 21.9 ± 1.7 | ns |
| Body weight change (g) | *** | 1.5 ± 0.9 | 1.5 ± 1.1 | ns | -5.6 ± 1.3 | -6.2 ± 1.3 | ns |
| Heart weight (mg) | *** | 134 ± 10 | 158 ± 13 | ns | 98 ± 7 | 126 ± 6 | ns |
| Heart weight (% of BW) | * | 0.49 ± 0.05 | 0.54 ± 0.05 | ns | 0.49 ± 0.05 | 0.58 ± 0.04 | * |
| Liver weight (g) | ** | 1.28 ± 0.15 | 1.44 ± 0.21 | ns | 0.84 ± 0.10 | 1.03 ± 0.20 | ns |
| Liver weight (% of BW) | ns | 4.6 ± 0.4 | 4.9 ± 0.4 | | 4.2 ± 0.4 | 4.7 ± 0.9 | |
| Epididymal white adipose (mg) | ** | 474 ± 92 | 569 ± 125 | ns | 181 ± 56 | 127 ± 27 | ns |
| Age at necropsy (days) | ns | 93 ± 7 | 95 ± 4 | | 91 ± 4 | 94 ± 5 | |
| *Metabolites* | | | | | | | |
| Glucose (mM) | * | 10.2 ± 1.9 | 10.5 ± 2.0 | ns | 7.5 ± 2.0 | 5.5 ± 3.2 | ns |
| Lactate (mM) | ns | 0.88 ± 0.37 | 0.82 ± 0.40 | | 1.47 ± 1.40 | 1.51 ± 0.68 | |
| Pyruvate (μM) | ns | 59 ± 29 | 49 ± 38 | | 44 ± 53 | 71 ± 52 | |
| β-hydroxybutyrate (μM) | ns | 42 ± 28 | 70 ± 44 | | 85 ± 46 | 204 ± 174 | |
| Free fatty acids (mM) | ns | 0.29 ± 0.03 | 0.32 ± 0.03 | | 0.32 ± 0.02 | 0.31 ± 0.09 | |
| Glycerol (mM) | ns | 0.20 ± 0.05 | 0.21 ± 0.04 | | 0.15 ± 0.03 | 0.23 ± 0.09 | |
| Triglycerides (mM) | * | 0.39 ± 0.14 | 0.44 ± 0.09 | ns | 0.19 ± 0.16 | 0.17 ± 0.07 | ns |
| *Amino acids* | | | | | | | |
| Phenylalanine (μM) | ns | 67 ± 24 | 52 ± 8 | | 67 ± 14 | 64 ± 18 | |
| Tyrosine (μM) | ns | 62 ± 23 | 44 ± 12 | | 48 ± 13 | 45 ± 15 | |
| Tryptophan (μM) | ** | 48 ± 12 | 44 ± 7 | ns | 74 ± 16 | 62 ± 12 | ns |
| Alanine (μM) | ns | 464 ± 137 | 353 ± 88 | | 388 ± 178 | 338 ± 103 | |
| Methionine (μM) | ** | 63 ± 13 | 56 ± 5 | ns | 47 ± 3 | 46 ± 4 | ns |
| Glycine (μM) | ns | 230 ± 50 | 201 ± 24 | | 185 ± 45 | 209 ± 45 | |
| Valine (μM) | ns | 174 ± 42 | 154 ± 25 | | 174 ± 46 | 159 ± 84 | |
| Leucine (μM) | ns | 95 ± 26 | 102 ± 18 | | 132 ± 40 | 114 ± 69 | |
| Isoleucine (μM) | ns | 82 ± 30 | 81 ± 26 | | 125 ± 41 | 96 ± 64 | |
| Glutamine (μM) | 0.051 | 749 ± 186 | 642 ± 105 | | 813 ± 64 | 699 ± 73 | |
| Asparagine (μM) | ns | 17 ± 6 | 15 ± 3 | | 15 ± 3 | 16 ± 3 | |
| Citrulline (μM) | ** | 48 ± 12 | 41 ± 5 | ns | 54 ± 8 | 65 ± 6 | ns |

| | | | | | | | |
|---|---|---|---|---|---|---|---|
| Ornithine (μM) | ns | 56 ± 15 | 65 ± 31 | | 49 ± 8 | 57 ± 12 | |
| Lysine (μM) | ns | 239 ± 57 | 204 ± 28 | | 208 ± 17 | 204 ± 31 | |
| Arginine (μM) | ns | 95 ± 26 | 69 ± 25 | | 68 ± 10 | 94 ± 22 | |
| Serine (μM) | ns | 117 ± 21 | 97 ± 19 | | 92 ± 14 | 91 ± 16 | |
| Proline (μM) | ns | 71 ± 25 | 54 ± 17 | | 48 ± 19 | 55 ± 12 | |
| Glutamate (μM) | ns | 52 ± 30 | 50 ± 24 | | 36 ± 14 | 29 ± 20 | |
| Aspartate (μM) | 0.067 | 8 ± 3 | 8 ± 1 | | 6 ± 1 | 6 ± 2 | |
| Total amino acids (mM) | ns | 2.73 ± 0.62 | 2.33 ± 0.23 | | 2.63 ± 0.31 | 2.45 ± 0.33 | |
| BCAA (μM) | ns | 350 ± 97 | 338 ± 62 | | 431 ± 125 | 368 ± 216 | |
| BCAA (%) | ns | 12.7 ± 1.1 | 14.5 ± 2.8 | | 16.4 ± 4.5 | 14.4 ± 6.5 | |
| BCAA/Ala (ratio) | ns | 0.76 ± 0.10 | 1.03 ± 0.44 | | 1.33 ± 0.73 | 1.29 ± 1.06 | |
| *Acylcarnitines* | | | | | | | |
| C0 (μM) | ns | 19.9 ± 5.2 | 17.9 ± 4.1 | ns | 22.9 ± 4.3 | 14.0 ± 7.3 | * |
| C2 (μM) | 0.070 | 4.52 ± 2.69 | 4.26 ± 1.87 | ns | 7.57 ± 2.44 | 3.73 ± 2.05 | * |
| C3 (μM) | ns | 0.17 ± 0.13 | 0.21 ± 0.20 | | 0.31 ± 0.25 | 0.23 ± 0.21 | |
| C4 (μM) | ns | 0.34 ± 0.28 | 0.36 ± 0.42 | | 0.50 ± 0.67 | 0.34 ± 0.36 | |
| C5 (μM) | ns | 0.07 ± 0.05 | 0.05 ± 0.05 | | 0.12 ± 0.13 | 0.08 ± 0.06 | |
| C4OH (μM) | * | 0.05 ± 0.03 | 0.04 ± 0.02 | ns | 0.12 ± 0.06 | 0.04 ± 0.02 | * |
| C6 (μM) | ns | 0.03 ± 0.02 | 0.03 ± 0.03 | | 0.04 ± 0.03 | 0.04 ± 0.03 | |
| C5OH (μM) | ns | 0.03 ± 0.03 | 0.02 ± 0.02 | | 0.04 ± 0.01 | 0.03 ± 0.02 | |
| C8 (μM) | ns | 0.02 ± 0.01 | 0.02 ± 0.01 | | 0.03 ± 0.01 | 0.03 ± 0.00 | |
| C3DC (μM) | ns | 0.01 ± 0.01 | 0.01 ± 0.00 | | 0.01 ± 0.01 | 0.01 ± 0.01 | |
| C4DC (μM) | ns | 0.04 ± 0.02 | 0.01 ± 0.02 | | 0.02 ± 0.02 | 0.01 ± 0.02 | |
| C12:1 (μM) | ** | 0.00 ± 0.01 | 0.09 ± 0.06 | ** | 0.01 ± 0.01 | 0.11 ± 0.10 | ns |
| C12 (μM) | ** | 0.02 ± 0.01 | 0.07 ± 0.04 | * | 0.02 ± 0.01 | 0.10 ± 0.06 | * |
| C6DC (μM) | ns | 0.01 ± 0.00 | 0.02 ± 0.01 | | 0.02 ± 0.01 | 0.03 ± 0.02 | |
| C14:2 (μM) | *** | 0.01 ± 0.01 | 0.18 ± 0.11 | ** | 0.01 ± 0.00 | 0.26 ± 0.18 | ** |
| C14:1 (μM) | *** | 0.03 ± 0.02 | 0.56 ± 0.24 | ** | 0.05 ± 0.03 | 0.77 ± 0.25 | * |
| C14 (μM) | ** | 0.03 ± 0.02 | 0.08 ± 0.03 | ns | 0.08 ± 0.03 | 0.12 ± 0.04 | ns |
| C8DC (μM) | ns | 0.00 ± 0.00 | 0.01 ± 0.01 | | 0.01 ± 0.00 | 0.01 ± 0.01 | |
| C16:1 (μM) | ns | 0.03 ± 0.01 | 0.05 ± 0.03 | | 0.06 ± 0.03 | 0.06 ± 0.03 | |
| C16 (μM) | * | 0.10 ± 0.05 | 0.16 ± 0.04 | ns | 0.21 ± 0.06 | 0.24 ± 0.10 | ns |
| C18:2 (μM) | * | 0.02 ± 0.01 | 0.03 ± 0.01 | ns | 0.04 ± 0.01 | 0.05 ± 0.02 | ns |
| C18:1 (μM) | ** | 0.06 ± 0.04 | 0.10 ± 0.04 | ns | 0.16 ± 0.05 | 0.15 ± 0.06 | ns |

| | | | | | | | |
|---|---|---|---|---|---|---|---|
| C18 (μM) | * | 0.03 ± 0.01 | 0.03 ± 0.01 | ns | 0.05 ± 0.01 | 0.04 ± 0.02 | ns |
| LCAC (μM) | *** | 0.30 ± 0.14 | 1.19 ± 0.44 | * | 0.66 ± 0.20 | 1.69 ± 0.47 | * |
| C14:1/C2 | *** | 0.01 ± 0.00 | 0.15 ± 0.10 | * | 0.01 ± 0.00 | 0.25 ± 0.14 | ** |

Values are the average ± SD. Groups sized are: WT (*Acadl*$^{+/+}$) chow n = 6, WT DR n = 6, LCAD KO (*Acadl*$^{-/-}$) chow n = 5, and LCAD KO DR n = 6. Statistical analysis of the data was performed using a nonparametric Kruskal-Wallis (KW) test with Dunn's multiple comparison post tests for WT versus KO. Significance (Sig.) is indicated as follows: ns = not significant, * < 0.05, ** < 0.01 and *** < 0.001.

**Table S2.** Analyzed variables in WT and LCAD KO mice after 4 weeks of ad libitum chow or dietary restriction. The effect of genotype is calculated in a post hoc test.

| | KW | WT | | post | LCAD KO | | post |
|---|---|---|---|---|---|---|---|
| Parameters | Sig. | Chow | DR | Sig. | Chow | DR | Sig. |
| Body weight (g) | *** | 27.7 ± 1.3 | 20.2 ± 1.6 | ** | 29.5 ± 2.6 | 21.9 ± 1.7 | * |
| Body weight change (g) | *** | 1.5 ± 0.9 | -5.6 ± 1.3 | * | 1.5 ± 1.1 | -6.2 ± 1.3 | ** |
| Heart weight (mg) | *** | 134 ± 10 | 98 ± 7 | * | 158 ± 13 | 126 ± 6 | * |
| Heart weight (% of BW) | * | 0.49 ± 0.05 | 0.49 ± 0.05 | ns | 0.54 ± 0.05 | 0.58 ± 0.04 | ns |
| Liver weight (g) | ** | 1.28 ± 0.15 | 0.84 ± 0.10 | ** | 1.44 ± 0.21 | 1.03 ± 0.20 | * |
| Liver weight (% of BW) | ns | 4.6 ± 0.4 | 4.2 ± 0.4 | | 4.9 ± 0.4 | 4.7 ± 0.9 | |
| Epididymal white adipose (mg) | ** | 474 ± 92 | 181 ± 56 | ns | 569 ± 125 | 127 ± 27 | ** |
| Age at necropsy (days) | ns | 93 ± 7 | 91 ± 4 | | 95 ± 4 | 94 ± 5 | |
| *Metabolites* | | | | | | | |
| Glucose (mM) | * | 10.2 ± 1.9 | 7.5 ± 2.0 | ns | 10.5 ± 2.0 | 5.5 ± 3.2 | * |
| Lactate (mM) | ns | 0.88 ± 0.37 | 1.47 ± 1.40 | | 0.82 ± 0.40 | 1.51 ± 0.68 | |
| Pyruvate (μM) | ns | 59 ± 29 | 44 ± 53 | | 49 ± 38 | 71 ± 52 | |
| β-hydroxybutyrate (μM) | ns | 42 ± 28 | 85 ± 46 | | 70 ± 44 | 204 ± 174 | |
| Free fatty acids (mM) | ns | 0.29 ± 0.03 | 0.32 ± 0.02 | | 0.32 ± 0.03 | 0.31 ± 0.09 | |
| Glycerol (mM) | ns | 0.20 ± 0.05 | 0.15 ± 0.03 | | 0.21 ± 0.04 | 0.23 ± 0.09 | |
| Triglycerides (mM) | * | 0.39 ± 0.14 | 0.19 ± 0.16 | ns | 0.44 ± 0.09 | 0.17 ± 0.07 | * |
| *Amino acids* | | | | | | | |
| Phenylalanine (μM) | ns | 67 ± 24 | 67 ± 14 | | 52 ± 8 | 64 ± 18 | |
| Tyrosine (μM) | ns | 62 ± 23 | 48 ± 13 | | 44 ± 12 | 45 ± 15 | |
| Tryptophan (μM) | ** | 48 ± 12 | 74 ± 16 | * | 44 ± 7 | 62 ± 12 | ns |
| Alanine (μM) | ns | 464 ± 137 | 388 ± 178 | | 353 ± 88 | 338 ± 103 | |
| Methionine (μM) | ** | 63 ± 13 | 47 ± 3 | ** | 56 ± 5 | 46 ± 4 | * |
| Glycine (μM) | ns | 230 ± 50 | 185 ± 45 | | 201 ± 24 | 209 ± 45 | |
| Valine (μM) | ns | 174 ± 42 | 174 ± 46 | | 154 ± 25 | 159 ± 84 | |
| Leucine (μM) | ns | 95 ± 26 | 132 ± 40 | | 102 ± 18 | 114 ± 69 | |
| Isoleucine (μM) | ns | 82 ± 30 | 125 ± 41 | | 81 ± 26 | 96 ± 64 | |
| Glutamine (μM) | 0.051 | 749 ± 186 | 813 ± 64 | | 642 ± 105 | 699 ± 73 | |
| Asparagine (μM) | ns | 17 ± 6 | 15 ± 3 | | 15 ± 3 | 16 ± 3 | |
| Citrulline (μM) | ** | 48 ± 12 | 54 ± 8 | ns | 41 ± 5 | 65 ± 6 | ** |
| Ornithine (μM) | ns | 56 ± 15 | 49 ± 8 | | 65 ± 31 | 57 ± 12 | |
| Lysine (μM) | ns | 239 ± 57 | 208 ± 17 | | 204 ± 28 | 204 ± 31 | |

| | | | | | | | |
|---|---|---|---|---|---|---|---|
| Arginine (μM) | ns | 95 ± 26 | 68 ± 10 | | 69 ± 25 | 94 ± 22 | |
| Serine (μM) | ns | 117 ± 21 | 92 ± 14 | | 97 ± 19 | 91 ± 16 | |
| Proline (μM) | ns | 71 ± 25 | 48 ± 19 | | 54 ± 17 | 55 ± 12 | |
| Glutamate (μM) | ns | 52 ± 30 | 36 ± 14 | | 50 ± 24 | 29 ± 20 | |
| Aspartate (μM) | 0.067 | 8 ± 3 | 6 ± 1 | ns | 8 ± 1 | 6 ± 2 | ns |
| Total amino acids (mM) | ns | 2.73 ± 0.62 | 2.63 ± 0.31 | | 2.33 ± 0.23 | 2.45 ± 0.33 | |
| BCAA (μM) | ns | 350 ± 97 | 431 ± 125 | | 338 ± 62 | 368 ± 216 | |
| BCAA (%) | ns | 12.7 ± 1.1 | 16.4 ± 4.5 | | 14.5 ± 2.8 | 14.4 ± 6.5 | |
| BCAA/Ala (ratio) | ns | 0.76 ± 0.10 | 1.33 ± 0.73 | | 1.03 ± 0.44 | 1.29 ± 1.06 | |
| *Acylcarnitines* | | | | | | | |
| C0 (μM) | ns | 19.9 ± 5.2 | 22.9 ± 4.3 | | 17.9 ± 4.1 | 14.0 ± 7.3 | |
| C2 (μM) | 0.070 | 4.52 ± 2.69 | 7.57 ± 2.44 | ns | 4.26 ± 1.87 | 3.73 ± 2.05 | ns |
| C3 (μM) | ns | 0.17 ± 0.13 | 0.31 ± 0.25 | | 0.21 ± 0.20 | 0.23 ± 0.21 | |
| C4 (μM) | ns | 0.34 ± 0.28 | 0.50 ± 0.67 | | 0.36 ± 0.42 | 0.34 ± 0.36 | |
| C5 (μM) | ns | 0.07 ± 0.05 | 0.12 ± 0.13 | | 0.05 ± 0.05 | 0.08 ± 0.06 | |
| C4OH (μM) | * | 0.05 ± 0.03 | 0.12 ± 0.06 | ns | 0.04 ± 0.02 | 0.04 ± 0.02 | ns |
| C6 (μM) | ns | 0.03 ± 0.02 | 0.04 ± 0.03 | | 0.03 ± 0.03 | 0.04 ± 0.03 | |
| C5OH (μM) | ns | 0.03 ± 0.03 | 0.04 ± 0.01 | | 0.02 ± 0.02 | 0.03 ± 0.02 | |
| C8 (μM) | ns | 0.02 ± 0.01 | 0.03 ± 0.01 | | 0.02 ± 0.01 | 0.03 ± 0.00 | |
| C3DC (μM) | ns | 0.01 ± 0.01 | 0.01 ± 0.01 | | 0.01 ± 0.00 | 0.01 ± 0.01 | |
| C4DC (μM) | ns | 0.04 ± 0.02 | 0.02 ± 0.02 | | 0.01 ± 0.02 | 0.01 ± 0.02 | |
| C12:1 (μM) | ** | 0.00 ± 0.01 | 0.01 ± 0.01 | ns | 0.09 ± 0.06 | 0.11 ± 0.10 | ns |
| C12 (μM) | ** | 0.02 ± 0.01 | 0.02 ± 0.01 | ns | 0.07 ± 0.04 | 0.10 ± 0.06 | ns |
| C6DC (μM) | ns | 0.01 ± 0.00 | 0.02 ± 0.01 | | 0.02 ± 0.01 | 0.03 ± 0.02 | |
| C14:2 (μM) | *** | 0.01 ± 0.01 | 0.01 ± 0.00 | ns | 0.18 ± 0.11 | 0.26 ± 0.18 | ns |
| C14:1 (μM) | *** | 0.03 ± 0.02 | 0.05 ± 0.03 | ns | 0.56 ± 0.24 | 0.77 ± 0.25 | ns |
| C14 (μM) | ** | 0.03 ± 0.02 | 0.08 ± 0.03 | ns | 0.08 ± 0.03 | 0.12 ± 0.04 | ns |
| C8DC (μM) | ns | 0.00 ± 0.00 | 0.01 ± 0.00 | | 0.01 ± 0.01 | 0.01 ± 0.01 | |
| C16:1 (μM) | ns | 0.03 ± 0.01 | 0.06 ± 0.03 | | 0.05 ± 0.03 | 0.06 ± 0.03 | |
| C16 (μM) | * | 0.10 ± 0.05 | 0.21 ± 0.06 | * | 0.16 ± 0.04 | 0.24 ± 0.10 | ns |
| C18:2 (μM) | * | 0.02 ± 0.01 | 0.04 ± 0.01 | ns | 0.03 ± 0.01 | 0.05 ± 0.02 | ns |
| C18:1 (μM) | ** | 0.06 ± 0.04 | 0.16 ± 0.05 | ** | 0.10 ± 0.04 | 0.15 ± 0.06 | ns |
| C18 (μM) | * | 0.03 ± 0.01 | 0.05 ± 0.01 | ** | 0.03 ± 0.01 | 0.04 ± 0.02 | ns |
| LCAC (μM) | *** | 0.30 ± 0.14 | 0.66 ± 0.20 | ns | 1.19 ± 0.44 | 1.69 ± 0.47 | ns |
| C14:1/C2 | *** | 0.01 ± 0.00 | 0.01 ± 0.00 | ns | 0.15 ± 0.10 | 0.25 ± 0.14 | ns |

Values are the average ± SD. Groups sized are: WT (*Acadl*$^{+/+}$) chow n = 6, WT DR n = 6, LCAD KO (*Acadl*$^{-/-}$) chow n = 5, and LCAD KO DR n = 6. Statistical analysis of the data was performed using a nonparametric Kruskal-Wallis (KW) test with Dunn's multiple comparison post tests for WT versus KO. Significance (Sig.) is indicated as follows: ns = not significant, * < 0.05, ** < 0.01 and *** < 0.001.

**Table S3.** Analyzed echocardiography variables in WT and LCAD KO mice at baseline (week 0).

| Parameter | t test | WT | LCAD KO |
|---|---|---|---|
| Heart rate (bpm) | | 495 ± 31 | 498 ± 29 |
| Body temperature (°C) | | 36.7 ± 0.7 | 37.1 ± 0.7 |
| Body weight (g) | ** | 25.7 ± 1.2 | 27.6 ± 1.4 |
| PWd (mm) | | 0.81 ± 0.16 | 0.91 ± 0.16 |
| PWs (mm) | | 1.03 ± 0.24 | 1.13 ± 0.22 |
| AWd (mm) | * | 0.89 ± 0.12 | 1.00 ± 0.09 |
| AWs (mm) | * | 1.25 ± 0.18 | 1.46 ± 0.20 |
| LVIDd (mm) | | 4.46 ± 0.16 | 4.38 ± 0.28 |
| LVIDs (mm) | | 3.29 ± 0.33 | 3.35 ± 0.46 |
| EF (%) | | 51.6 ± 8.1 | 50.6 ± 8.0 |
| FS (%) | | 26.4 ± 5.0 | 25.8 ± 5.0 |

Values are the average ± SD. Groups sized are: WT ($Acadl^{+/+}$) n = 12 and LCAD KO ($Acadl^{-/-}$) n = 11. Statistical analysis of the data was performed using a t test (two-tailed, two-sample equal variance). Significance (Sig.) is indicated as follows: * < 0.05 and ** < 0.01.

**Table S4.** Analyzed echocardiography variables in WT and LCAD KO mice after 4 weeks ad libitum chow or dietary restriction. The effect of the dietary intervention is calculated in a post hoc test.

| | KW | Chow | | post | Dietary restriction | | post |
|---|---|---|---|---|---|---|---|
| Parameter | Sig. | WT | LCAD KO | Sig. | WT | LCAD KO | Sig. |
| Heart rate (bpm) | *** | 516 ± 16 | 500 ± 32 | ns | 413 ± 35 | 395 ± 32 | ns |
| Body temperature (°C) | ns | 36.8 ± 0.2 | 36.9 ± 0.8 | | 36.8 ± 0.6 | 36.2 ± 0.9 | |
| Body weight (g) | *** | 28.6 ± 1.2 | 29.9 ± 2.1 | ns | 20.6 ± 1.0 | 22.0 ± 1.1 | ns |
| PWd (mm) | * | 0.69 ± 0.16 | 0.83 ± 0.04 | ns | 0.52 ± 0.14 | 0.86 ± 0.35 | * |
| PWs (mm) | ns | 0.92 ± 0.18 | 1.04 ± 0.17 | | 0.73 ± 0.22 | 1.08 ± 0.35 | |
| AWd (mm) | * | 0.96 ± 0.09 | 1.01 ± 0.04 | ns | 0.76 ± 0.13 | 0.97 ± 0.17 | * |
| AWs (mm) | ns | 1.32 ± 0.22 | 1.32 ± 0.08 | | 1.10 ± 0.23 | 1.22 ± 0.14 | |
| LVIDd (mm) | ns | 4.50 ± 0.21 | 4.63 ± 0.47 | | 4.44 ± 0.29 | 4.49 ± 0.39 | |
| LVIDs (mm) | ns | 3.18 ± 0.28 | 3.52 ± 0.50 | | 3.27 ± 0.37 | 3.56 ± 0.31 | |
| EF (%) | * | 56.3 ± 5.0 | 47.8 ± 6.7 | ns | 51.9 ± 6.8 | 42.1 ± 5.4 | * |
| FS (%) | * | 29.4 ± 3.2 | 24.1 ± 3.9 | ns | 26.6 ± 4.4 | 20.7 ± 3.1 | ns |

Values are the average ± SD. Groups sized are: WT (*Acadl*$^{+/+}$) chow n = 6, WT DR n = 6, LCAD KO (*Acadl*$^{-/-}$) chow n = 5, and LCAD KO DR n = 6. Statistical analysis of the data was performed using a nonparametric Kruskal-Wallis (KW) test with Dunn's multiple comparison post tests for WT versus KO. Significance (Sig.) is indicated as follows: ns = not significant, * < 0.05, ** < 0.01 and *** < 0.001.

**Table S5.** Analyzed echocardiography variables in WT and LCAD KO mice after 4 weeks ad libitum chow or dietary restriction. The effect of genotype is calculated in a post hoc test.

| Parameter | KW Sig. | WT Chow | WT DR | post Sig. | LCAD KO Chow | LCAD KO DR | post Sig. |
|---|---|---|---|---|---|---|---|
| Heart rate (bpm) | *** | 516 ± 16 | 413 ± 35 | ** | 500 ± 32 | 395 ± 32 | * |
| Body temperature (°C) | ns | 36.8 ± 0.2 | 36.8 ± 0.6 | | 36.9 ± 0.8 | 36.2 ± 0.9 | |
| Body weight (g) | *** | 28.6 ± 1.2 | 20.6 ± 1.0 | ** | 29.9 ± 2.1 | 22.0 ± 1.1 | * |
| PWd (mm) | * | 0.69 ± 0.16 | 0.52 ± 0.14 | ns | 0.83 ± 0.04 | 0.86 ± 0.35 | ns |
| PWs (mm) | ns | 0.92 ± 0.18 | 0.73 ± 0.22 | | 1.04 ± 0.17 | 1.08 ± 0.35 | |
| AWd (mm) | * | 0.96 ± 0.09 | 0.76 ± 0.13 | ns | 1.01 ± 0.04 | 0.97 ± 0.17 | ns |
| AWs (mm) | ns | 1.32 ± 0.22 | 1.10 ± 0.23 | | 1.32 ± 0.08 | 1.22 ± 0.14 | |
| LVIDd (mm) | ns | 4.50 ± 0.21 | 4.44 ± 0.29 | | 4.63 ± 0.47 | 4.49 ± 0.39 | |
| LVIDs (mm) | ns | 3.18 ± 0.28 | 3.27 ± 0.37 | | 3.52 ± 0.50 | 3.56 ± 0.31 | |
| EF (%) | * | 56.3 ± 5.0 | 51.9 ± 6.8 | ns | 47.8 ± 6.7 | 42.1 ± 5.4 | ns |
| FS (%) | * | 29.4 ± 3.2 | 26.6 ± 4.4 | ns | 24.1 ± 3.9 | 20.7 ± 3.1 | ns |

Values are the average ± SD. Groups sized are: WT (*Acadl*$^{+/+}$) chow n = 6, WT DR n = 6, LCAD KO (*Acadl*$^{-/-}$) chow n = 5, and LCAD KO DR n = 6. Statistical analysis of the data was performed using a nonparametric Kruskal-Wallis (KW) test with Dunn's multiple comparison post tests for WT versus KO. Significance (Sig.) is indicated as follows: ns = not significant, * < 0.05, ** < 0.01 and *** < 0.001.

### Figure S1

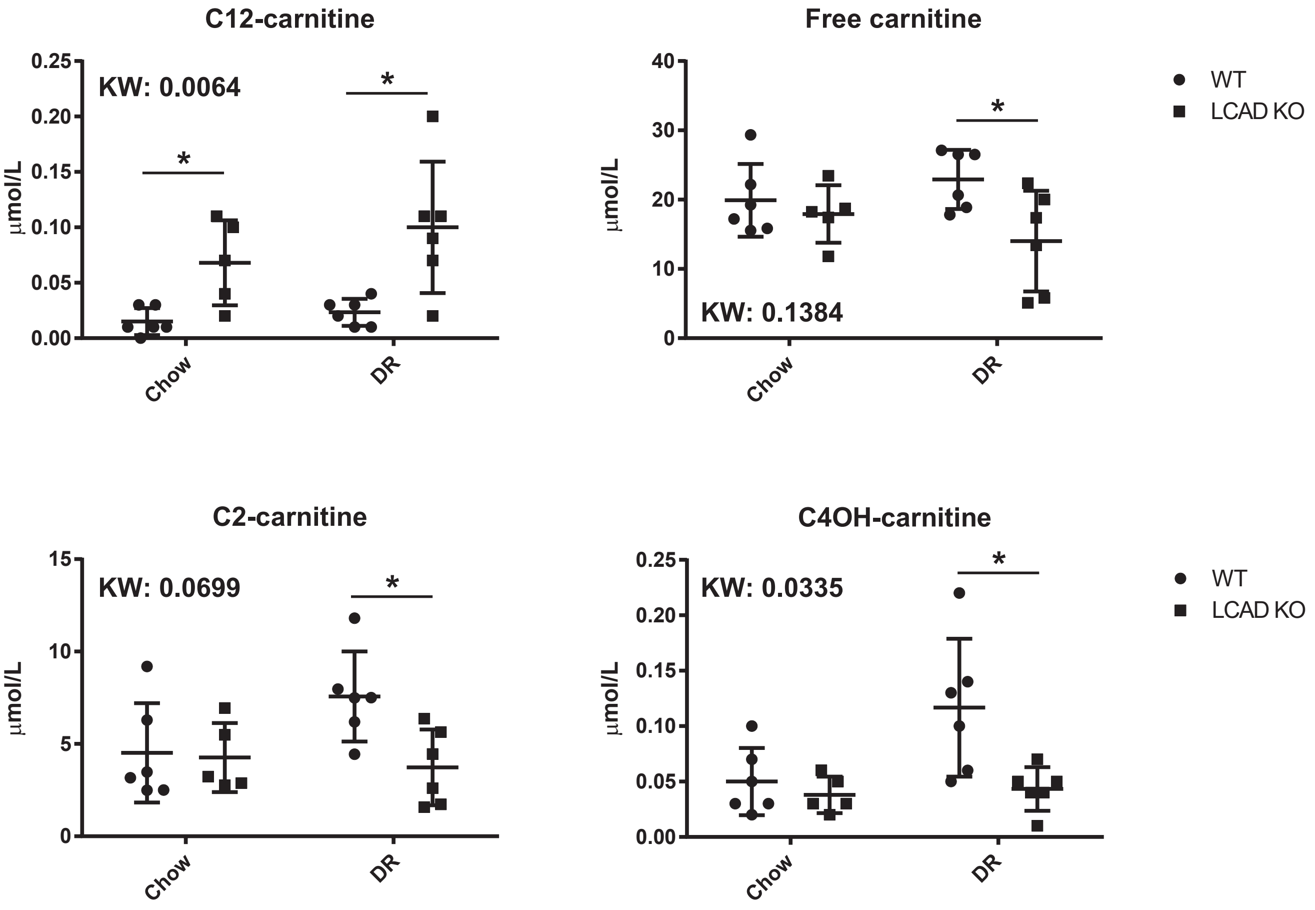
C12-carnitine
KW: 0.0064
μmol/L
0.25
0.20
0.15
0.10
0.05
0.00
*
*
Chow
DR
Free carnitine
KW: 0.1384
μmol/L
40
30
20
10
0
*
Chow
DR
WT
LCAD KO
C2-carnitine
KW: 0.0699
μmol/L
15
10
5
0
*
Chow
DR
C4OH-carnitine
KW: 0.0335
μmol/L
0.25
0.20
0.15
0.10
0.05
0.00
*
Chow
DR
WT
LCAD KO

**Figure S2**

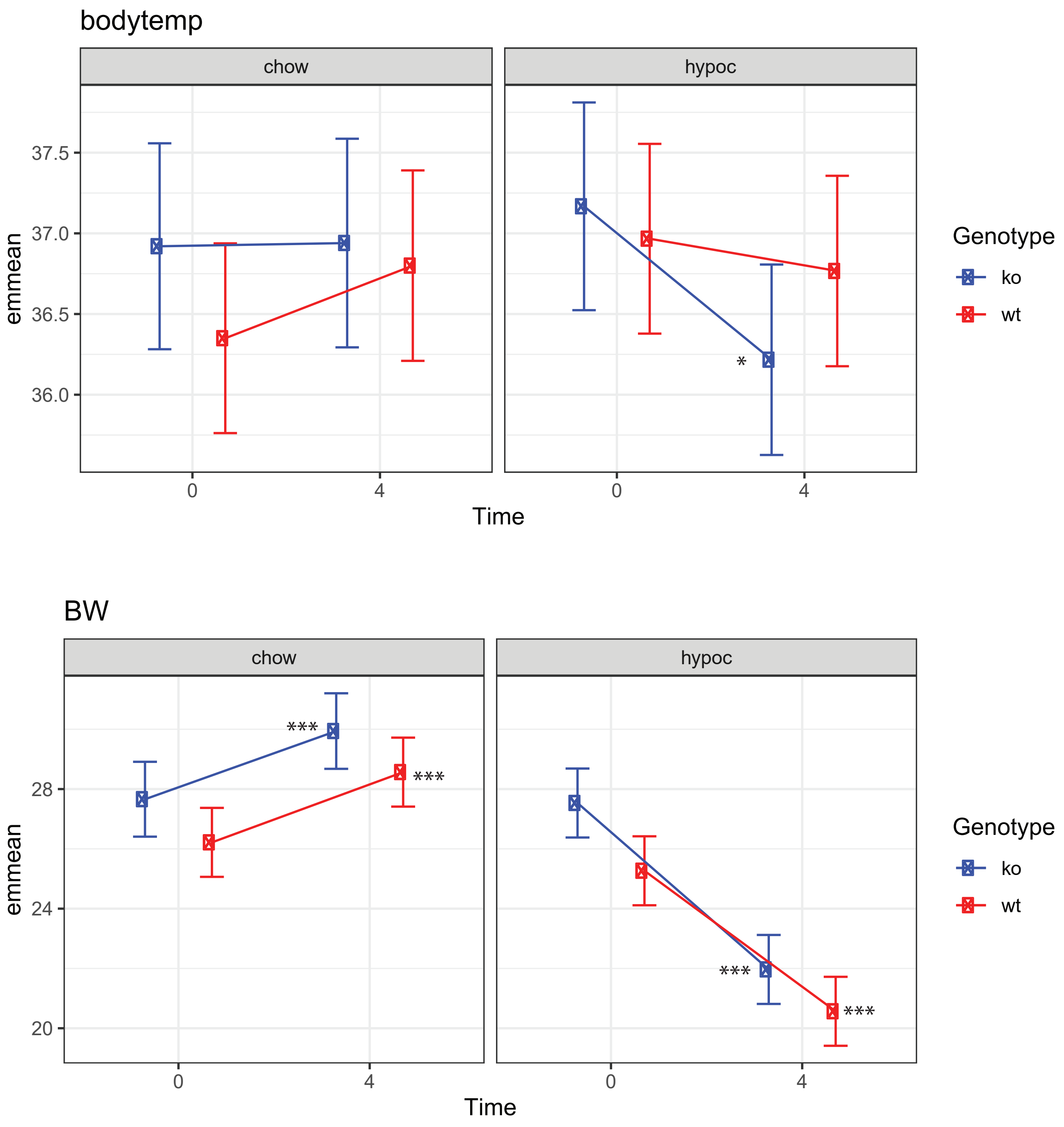
bodytemp
chow
hypoc
37.5
37.0
36.5
36.0
emmean
0
4
Time
*
Genotype
ko
wt
BW
chow
hypoc
28
24
20
emmean
***
***
***
***
0
4
Time
Genotype
ko
wt
